# Phase Separation of EMB1579 Drives P-TEFb-Mediated Phosphorylation of RNA Pol II CTD in Arabidopsis

**DOI:** 10.64898/2026.11.25.769200

**Authors:** Xuan Zhang, Yuqin Ding, Yiling Zhang, Ping Han, Shanjin Huang

## Abstract

Phosphorylation of the C-terminal domain (CTD) of RNA polymerase II (Pol II) by the cyclin-dependent kinase C (CDKC)-CYCT complex is crucial for Pol II-mediated transcription in Arabidopsis. Here, we show that phase separation of the intrinsically disordered protein EMBRYO DEFECTIVE 1579 (EMB1579) orchestrates this process. CDKC;2, the major CDKC family protein in Arabidopsis, interacts with EMB1579 at least in part through its C-terminal intrinsically disordered region (IDR), and the two proteins co-condense in vitro, with EMB1579 acting as the primary driver of condensate formation. In vivo, CDKC;2 colocalizes with EMB1579 condensates, and EMB1579 phase separation enhances CDKC;2-mediated CTD phosphorylation both in vitro and in planta. Strikingly, EMB1579 condensates recruit not only CDKC;2 but also CYCT and synthetic CTD peptides, functioning as reaction hubs for Pol II CTD phosphorylation. This study reveals a novel mechanism by which phase-separated biomolecular condensates spatially organize transcriptional kinases to precisely regulate Pol II activity, advancing our understanding of eukaryotic transcription regulation.

## Introduction

The regulation of RNA polymerase II (RNA Pol II)-mediated transcription is central to plant growth, development, and environmental adaptation (Obermeyer et al. 2024; Grasser 2025). A key structural feature of RNA Pol II is its carboxyl-terminal domain (CTD), composed of repetitive heptapeptide sequences (YSPTSPS) conserved across eukaryotes (Zaborowska, Egloff, and Murphy 2016; Coudreuse et al. 2010). Although dispensable for catalytic activity, the CTD is indispensable for coordinating multiple steps of transcription and co-transcriptional processes, including mRNA processing, chromatin remodeling, and transcriptional elongation (Bird, Zorio, and Bentley 2004; Corden 2013; Coudreuse et al. 2010; Jeronimo, Bataille, and Robert 2013). The CTD undergoes dynamic post-translational modification, including phosphorylation, acetylation, methylation, ubiquitination, and O-GlcNAcylation (Heidemann et al. 2013; Lu et al. 2016; Ranuncolo et al. 2012; Li et al. 2007; Schroder et al. 2013; Dias et al. 2015), which modulates its interactions with regulatory proteins. Among these, phosphorylation is particularly well-characterized, as it dictates the recruitment of specific protein complexes during different transcriptional phases, thereby influencing transcription at multiple levels.

Cyclin-dependent kinases (CDKs) are critical regulators of CTD phosphorylation. In *Arabidopsis thaliana*, 30 CDK and CDK-like (CDK-L) genes are classified into eight subgroups (CDKA to CDKG and CDK-L). While CDKA and the plant-specific CDKB are primarily associated with cell cycle control, CDKC, CDKD, CDKE, and CDKG have emerged as key players in transcriptional regulation (Barroco et al. 2003; Hajheidari, Koncz, and Eick 2013; Zheng 2022). For example, mammalian Cdk9/CycT complexes, components of the positive transcription elongation factor b (P-TEFb), phosphorylate CTD Serine 2 (Ser2) to promote transcriptional elongation. In Arabidopsis, CDKC proteins, orthologs of Cdk9, interact with CYCT cyclins to globally regulate Ser2 phosphorylation levels (Barroco et al. 2003; Wang et al. 2014; Uehara et al. 2022). The CDKC family comprises two members, CDKC;1 and CDKC;2 (Barroco et al. 2003), with CDKC;2 exhibiting higher expression across tissues and playing roles in lateral organ development, circadian clock regulation, flowering time, and stress responses (Cui et al. 2007; Li et al. 2014; Wang et al. 2014; Zhao et al. 2017a; Uehara et al. 2022). Despite these insights, the regulatory mechanisms governing CDKC;2’s function in CTD phosphorylation remain poorly understood.

Recent advances highlight the role of non-membranous biomolecular condensates, formed via liquid-liquid phase separation (LLPS) (Alberti and Hyman 2021; Zhang, Mehta, and Zhang 2021; Shin and Brangwynne 2017), in cellular organization. LLPS-driven condensates are now recognized as key regulators of transcriptional processes, including chromatin remodeling, signal transduction, and environmental response (Wang and Liu 2019; Chen et al. 2022; Sanulli et al. 2019; Li et al. 2012; Jung et al. 2020). In plants, LLPS mechanisms have been implicated in controlling transcriptional events such as RNA 3’ end processing, pre-mRNA splicing, and alternative polyadenylation (Simpson et al. 2003; Wu et al. 2016; Miao et al. 2026; Kyung, Jeon, and Lee 2022; Wu et al. 2020). For instance, the floral repressor FLOWERING LOCUS C (FLC) is regulated by LLPS-mediated compartmentalization, in which FLL2 promotes the formation of FCA nuclear bodies that concentrate 3’ end processing factors to enhance polyadenylation at the *FLC* locus (Fang et al. 2019). Similarly, phase separation of HRLP (hnRNP R-like protein) with splicing factors facilitates R-loop formation and RNA Pol II recruitment, influencing *FLC* splicing (Zhang et al. 2022). Notably, the Arabidopsis protein EMBRYO DEFECTIVE 1579 (EMB1579) undergoes LLPS to regulate global transcription and splicing, including *FLC* expression (Zhang et al. 2020b). Recently, it was shown that EMB1579 maintains root meristem stem cell homeostasis through the coordinated action of transcriptional regulation and RNA splicing, with its function dependent on physical interaction with splicing factors (Aljedaani et al. 2025). However, the molecular principles governing how EMB1579 controls transcription remain elusive.

Here, we report that EMB1579 physically interacts with CDKC;2 to enhance its kinase activity toward RNA Pol II CTD Ser2 phosphorylation. Loss of EMB1579 reduces Pol II Ser2 phosphorylation *in vivo*. We further demonstrate that CDKC;2 is recruited into EMB1579 condensates to form larger, more dynamic assemblies, and that EMB1579 mutants defective in LLPS exhibit reduced capacity to potentiate CDKC;2 activity. Our findings elucidate a novel mechanism linking protein LLPS to the regulation of Pol II CTD phosphorylation and advance understanding of how liquid condensates govern transcriptional control in plants.

## Results

### EMB1579 Mutants Phenocopy *cdkc;2* Mutants in Transcription and Plant Development

Our previous study demonstrated that EMB1579 regulates global transcription and splicing (Zhang et al. 2020b). To understand how EMB1579 functions in transcriptional regulation, we analyzed proteins that potentially interact with EMB1579, focusing on those implicated in RNA Pol II-mediated transcription (Figure S1A; (Zhang et al. 2020b)). Of particular interest was the CDKC family proteins (CDKC;1 and CDKC;2) (Figure S1B), the primary RNA Pol II kinases regulating transcription. The physical interaction between EMB1579 and CDKC kinases has been previously verified (Zhang et al. 2020b). We focused on *CDKC;2* because in silico gene expression analyses revealed that its expression is substantially higher than that of *CDKC;1* throughout development (Figure S2), and previous studies in this field have primarily investigated the functional roles of CDKC;2 (Kitsios et al. 2008; Wang et al. 2014; Cui et al. 2007).

Phenotypic analyses first confirmed that the *cdkc;2* mutant exhibits a late-flowering phenotype (Figure S3), which is fully consistent with findings from previous independent studies (Cui et al. 2007; Wang et al. 2014). Notably, side-by-side phenotypic characterization revealed substantial overlap in developmental defects between *cdkc;2* and *emb1579* mutants. Both mutants display shortened primary roots (Figure 1A, B), delayed flowering time, elevated transcript levels of the floral repressor *FLC* (Figure 1C-E; (Zhang et al. 2020b; Cui et al. 2007)), and enlarged rosette leaves (Figure 1F-H), even though *emb1579* consistently exhibits more severe phenotype. Given the well-documented regulatory axis in which loss of CDKC;2 globally reduces RNA Polymerase II Ser2 phosphorylation, directly suppresses transcription of the *FLC* antisense transcript COOLAIR, and consequently leads to upregulated *FLC* expression **(Wang et al. 2014)**, we hypothesized that the extensive phenotypic and molecular overlap between *cdkc;2* and *emb1579* arises from a functional link where EMB1579 participates in the CDKC;2-mediated transcriptional regulation pathway that controls Pol II phosphorylation. This proposed regulatory connection is further corroborated by our parallel gene expression assays: the rapid transcriptional induction of the stress-responsive marker genes *RAB28* and *HSP70* (Hundertmark and Hincha 2008; Borrell et al. 2002) is significantly impaired in both *emb1579* and *cdkc;2* mutants under the same treatment conditions (Figure 1I, J). Collectively, these convergent lines of genetic and molecular evidence support the conclusion that EMB1579 modulates multiple aspects of plant growth and development, at least in part, through functionally regulating the activity of the CDKC;2-dependent transcriptional regulatory module.

**Figure 1.**
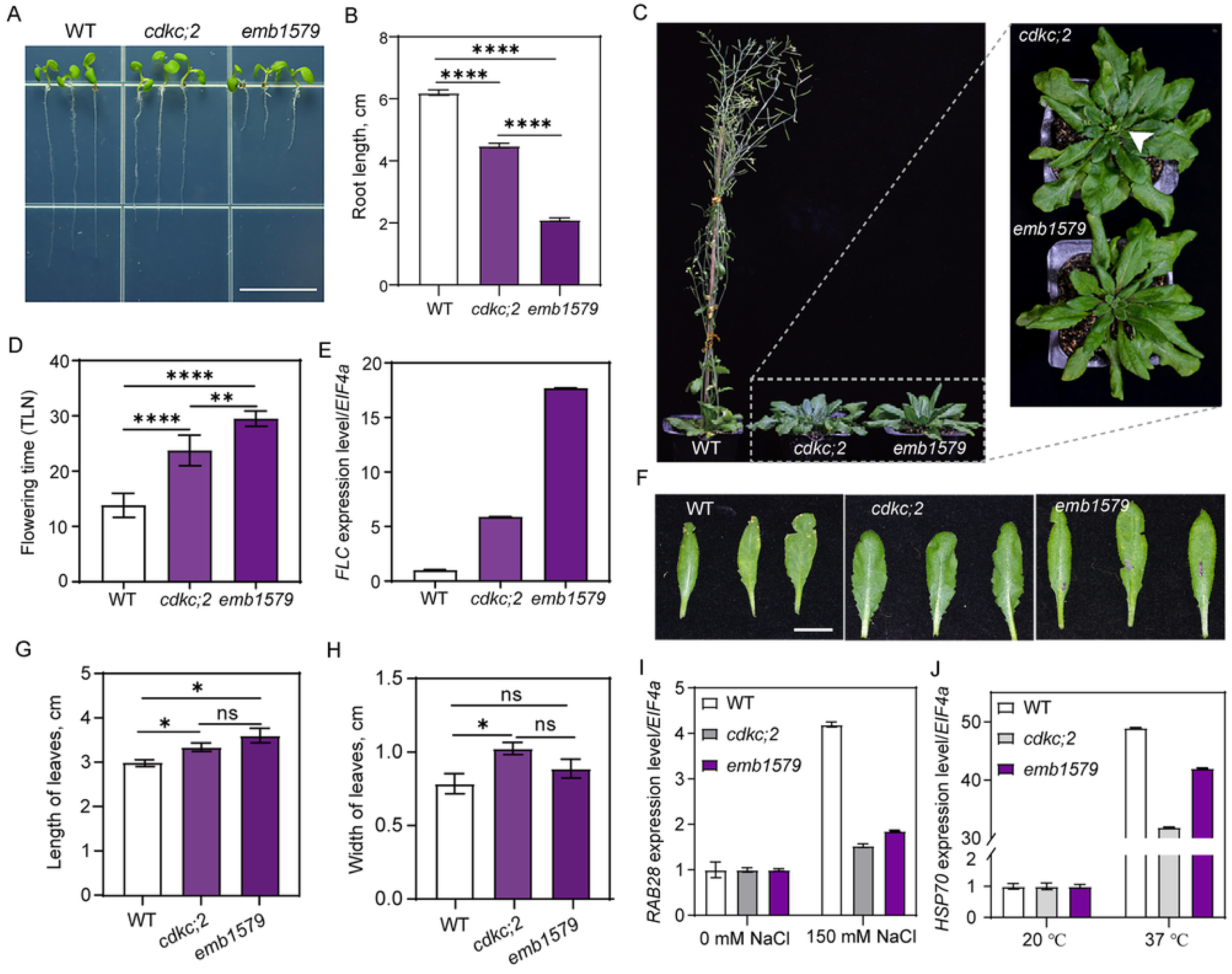
The *emb1579* mutant phenocopies *cdkc;2* mutants. **(A)** Images of wild type, *cdkc;2* and *emb1579* seedlings that were grown for 7 days on 1/2 MS medium. Scale bars, 1 cm. **(B)** Length of root from the seedlings shown in **(B).** The data are mean ± SEM; n = 50. Statistical comparison was performed with two-tailed paired Student’s *t*-test, ***P<0.001. **(C)** Images of wild type, *cdkc;2* and *emb1579* plants after 60 days of normal growth in soil under long light conditions. Front view (left panel) and vertical view (right panel) of corresponding plants. The white arrow in the bottom panel indicate the flower bud of *cdkc;2*. **(D)** Flowering time (total leaf number, TLN). Total number of rosette leaves were counted when the plants first flowers bloomed. The data are means ± SEM; n=20. Statistical comparison was performed with two-tailed paired Student’s *t*-test, **P<0.01 and ***P<0.001. **(E)** *FLC* expression levels in 7-day-old seedlings in different genetic backgrounds normalized to *EIF4a* expression are shown relative to the *FLC* expression level in wild type set at 1.0. The data are means ± SEM; n = 3. Statistical comparison was performed with two-tailed paired Student’s *t*-test, *P<0.05. ns, not significant. **(F)** Images of leaf phenotype of wild type, *cdkc2* and *emb1579*. The rosette leaves of plants grown for 8 weeks were selected and photographed for recording. Scale bars, 1 cm. **(G**, **H)** The length and maximum width statistics of the maximum rosette leaves of the wild type, *cdkc2* and *emb1579* grown for 8 weeks. The data are means ± SEM, n=10. Statistical comparison was performed with two-tailed paired Student’s *t*-test, *P<0.05. ns, not significant. (**I**, **J**) *RAB28* and *HSP70* expression level in 7-day-old seedlings in different genetic backgrounds with or without stresses stimulation. *RAB28* and *HSP70* expression levels normalized to *EIF4a* expression. The gene expression levels in plants without stresses treatment are set as 1.0. The data are means ± SEM; n = 3.

### EMB1579 Enhances CDKC;2 Kinase Activity to Regulate Pol II CTD Phosphorylation and *FLC* Transcription

To determine whether EMB1579 regulates development through CDKC;2-mediated Pol II CTD phosphorylation, we first examined Pol II CTD phosphorylation levels *in vivo*. Both *cdkc;2* and *emb1579* mutants exhibited reduced Pol II CTD Ser2 phosphorylation, with a more pronounced reduction in *cdkc;2* than in *emb1579* (Figure 2A), suggesting that EMB1579 positively regulates CDKC;2 kinase activity. Consistent with this, *in vitro* kinase assays revealed that EMB1579 enhances CDKC;2-mediated phosphorylation of the Pol II CTD Ser5 residue in a dose-dependent manner (Figure 2B, C).

**Figure 2.**
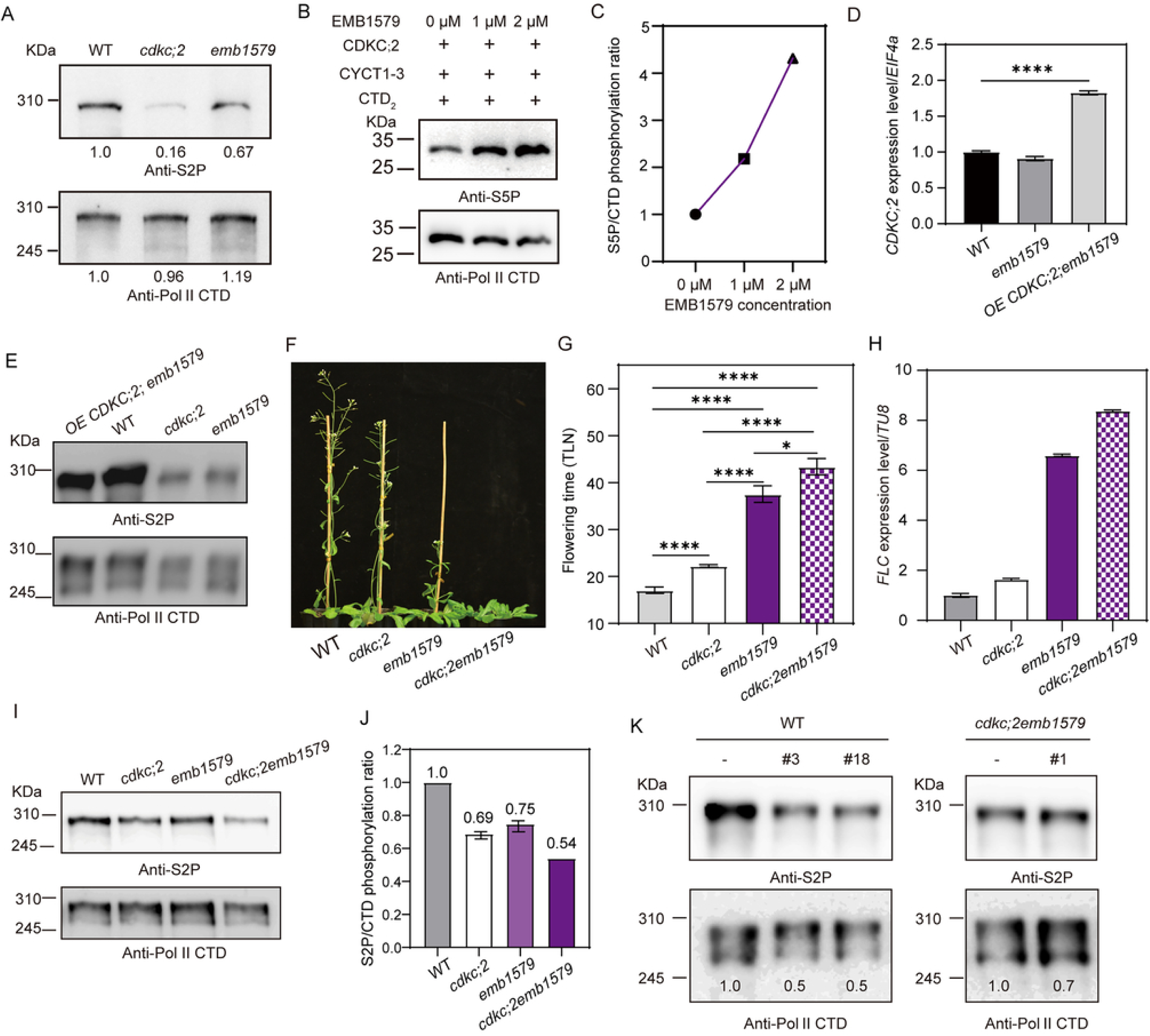
The level of pol II CTD phosphorylation is reduced in *emb1579* mutant and EMB1579 acts upstream of CDKC;2. **(A)** RNA pol II Ser2 phosphorylation ratio in different genetic backgrounds. Western blots of protein extracted from 12-day-old seedlings. Membranes were probed with antibodies against CTD Ser2 P Pol II (3E10) or total Pol II (8WG16). **(B)** In vitro kinase assays showing dose-dependent enhancement of CDKC;2 activity by EMB1579. Purified EGFP-CDKC;2, CYCT1-3-YFP, and GST-CTD2 were incubated with increasing concentrations of EMB1579 in phosphorylation buffer. Membranes were probed with antibodies against Ser5-P Pol II (D9N5I) or total Pol II (8WG16). **(C)** Quantification of relative amount of CTD phosphorylation. S5P/CTD was determined to assess the relative activity of CDKC;2/CYCT1-3 in phosphorylating pol II CTD in the presence of different concentrations of EMB1579. **(D, E)** Overexpression of CDKC;2 partially rescues Pol II phosphorylation in *emb1579*. **(D)** Relative CDKC;2 expression levels. CDKC;2 was overexpressed in *emb1579* mutant, designated as OE CDKC;2 *emb1579*. *EIF4A* was used as the loading control. RNA was extracted from 7-day-old wild-type, *cdkc;2*, *emb1579*, and OE CDKC;2; *emb1579* seedlings and analyzed by qRT-PCR. Transcript levels were normalized to *EIF4A* and set to 1 in wild-type. Data shown as mean ± SD (n = 3). Statistical comparison between wild type seedlings and the *emb1579* plants overexpressing *CDKC;2* was performed with two-tailed paired Student’s *t*-test, \*\*\*\**P* ≤ 0.0001. **(E)** Effect of *CDKC;2* overexpression on Pol II Ser2 phosphorylation in the *emb1579* background. Total protein was extracted from 7-day-old seedlings of the indicated genotypes and analyzed by immunoblotting. Blots were probed with S2P (3E10) to detect RNA Pol II Ser2 phosphorylation and with Pol II CTD (8WG16) to detect total RNA Pol II CTD. **(F)** Plant phenotypes of wild type, *cdkc;2*, *emb1579* and *cdkc;2emb1579* plants after 40 days of normal growth in soil under long light conditions. **(G)** Total number of rosette leaves of plants shown in (**F**) were counted when the plants first flower bloomed to assess the flowering time. The data are means ± SEM; n=20. Asterisks indicate significant differences between wild type seedlings and the other genotypes (two-tailed paired Student’s *t* test, \**P* ≤ 0.05; \*\*\*\**P* ≤ 0.0001). **(H)** *FLC* expression level of plants shown in (**F**) was plotted. *FLC* expression levels normalized to *TUB8* expression are shown relative to the *FLC* expression level in wild type set at 1.0. The data are means ± SEM; n = 3. **(I)** RNA pol II Ser2 phosphorylation level detection as that shown in (**A**). **(J)** RNA pol II Ser2 phosphorylation ratio in different genetic backgrounds. Ser2 phosphorylation level are shown relative to that in wild type set at 1.0. The data are means ± SEM; n = 7. **(K)** RNA pol II Ser2 phosphorylation ratio in wild type or *emb1579* when the expression level of CDKC;1 were knockdown. Ser2 phosphorylation level are shown relative to that in wild type or *cdkc;2emb1579* set at 1.0.

To confirm that the reduced pol II CTD phosphorylation in *emb1579* results specifically from impaired EMB1579-mediated enhancement of CDKC;2 activity, we overexpressed *CDKC;2* in the *emb1579* background (Figure 2D). Pol II CTD phosphorylation levels increased substantially in *emb1579* plants overexpressing *CDKC;2*, reaching levels comparable to wild type (Figure 2E). To further investigate the genetic relationship between EMB1579 and CDKC;2 *in vivo*, we generated *cdkc;2 emb1579* double mutants. These plants exhibited a significant flowering delay relative to either single mutant (Figure 2F, G), correlating with substantially elevated *FLC* transcript levels (Figure 2H). Notably, Pol II CTD phosphorylation was further reduced in the double mutant compared with the *cdkc;2* single mutant (Figure 2I, J), indicating that EMB1579 positively regulates additional targets beyond CDKC;2 in Pol II CTD phosphorylation.

To determine whether loss of EMB1579-mediated enhancement of CDKC;1 activity contributes to the further reduction in Pol II CTD phosphorylation observed in *cdkc;2 emb1579* double mutants, we silenced *CDKC;1* via RNAi in both wild-type and *cdkc;2 emb1579* backgrounds, as simultaneous loss of both redundant *CDKC* genes is lethal. CDKC;1 depletion caused a marked decline in Pol II CTD phosphorylation in wild type, comparable to that seen in *cdkc;2* single mutants (Figure 2K), confirming that CDKC;1 functions in maintaining Pol II CTD phosphorylation levels *in vivo*. Importantly, CDKC;1 loss further reduced CTD phosphorylation in the *cdkc;2 emb1579* background (Figure 2K), indicating that the loss of EMB1579-mediated enhancement of CDKC;1 activity accounts for the additional drop in Pol II CTD phosphorylation in the double mutant relative to *cdkc;2* alone. Together, these results establish that EMB1579 positively regulates both CDKC;1 and CDKC;2 to sustain Pol II CTD phosphorylation.

To further probe the functional relationship between EMB1579 and CDKC kinases, we treated 7-day-old wild-type and *emb1579* seedlings with the CDKC inhibitor BML259. Pol II CTD phosphorylation decreased in a dose-dependent manner in both genotypes (Figure S4A, B), yet at every concentration the reduction was more severe in *emb1579* than in wild type (Figure S4A, B). This heightened sensitivity is consistent with the loss of EMB1579-mediated kinase enhancement: when CDKC activity is already compromised by the absence of EMB1579, pharmacological inhibition produces a proportionally greater effect. Correspondingly, *FLC* transcription increased with BML259 treatment in both backgrounds (Figure S4C). Together, these pharmacological data support a model in which EMB1579 acts upstream of CDKC;2 to enhance Pol II CTD phosphorylation, thereby restraining *FLC* expression presumably by promoting the transcription of the *FLC* antisense transcript COOLAIR and promoting normal plant growth and development.

### CDKC;2 Colocalizes with EMB1579 Condensates and Co-condenses with EMB1579 *In Vitro*

To examine how CDKC;2 and EMB1579 function coordinately *in vivo*, we generated functional EMB1579-3×GFP and tdTomato-CDKC;2 fusion proteins stably expressed in the *cdkc;2 emb1579* double mutant background (Figure 3A). Both proteins localized to the nucleus and formed distinct foci in Arabidopsis root cells, as previously reported (Zhang et al. 2020b; Kitsios et al. 2008). Notably, CDKC;2 foci partially colocalized with EMB1579 condensates (Figure 3A). Given the previously demonstrated physical interaction between EMB1579 and CDKC;2 (Zhang et al. 2020b), we hypothesized that the two proteins co-condense through this interaction.

**Figure 3.**
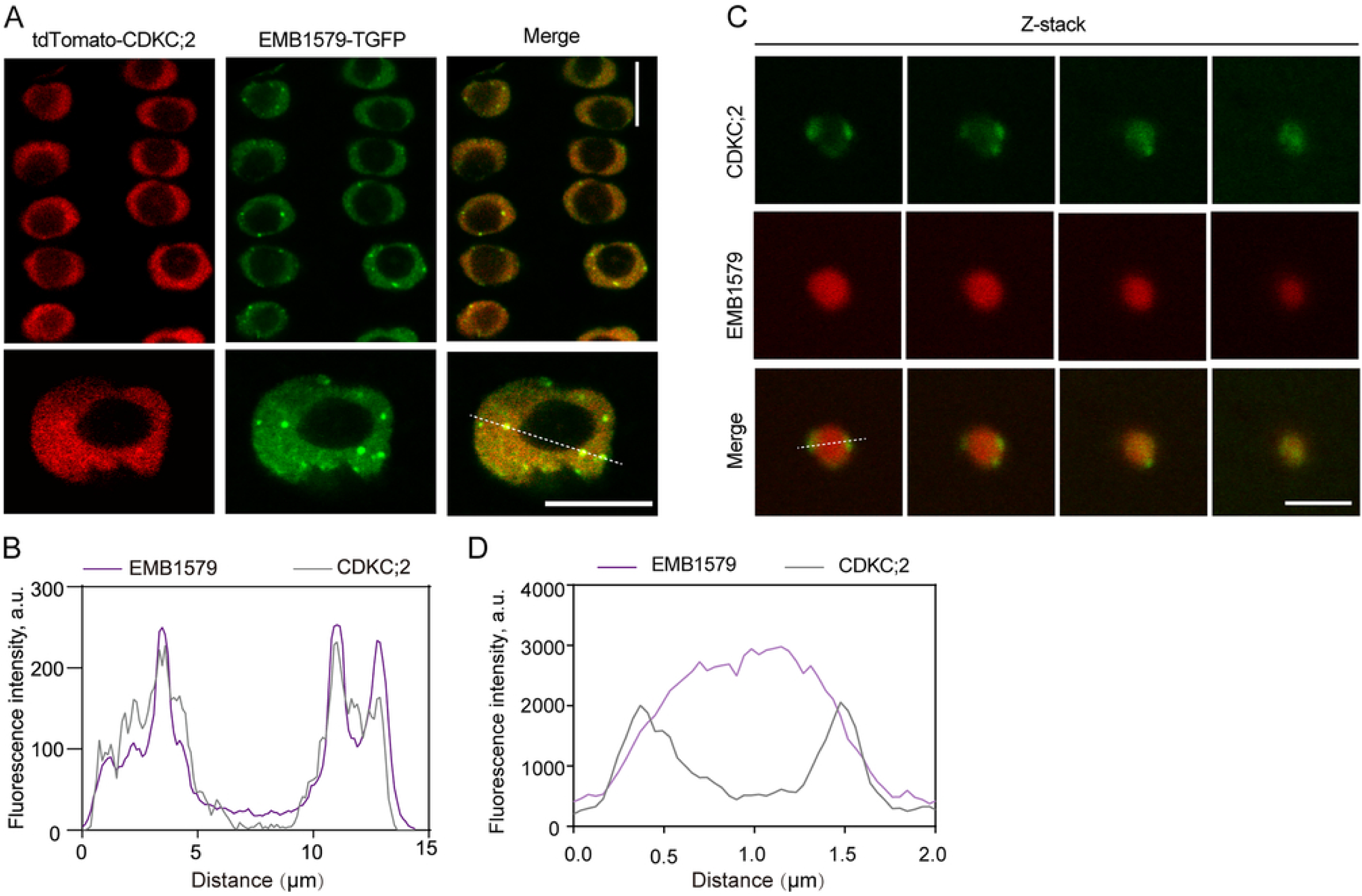
EMB1579 colocalizes with CDKC;2 in the nucleus and co-condenses with CDKC;2 *in vitro*. **(A)** Representative confocal images of CDKC;2 and EMB1579 condensates in Arabidopsis root cells. White dashed lines indicate regions of co-localization. Scale bars, 5 μm. **(B)** Line-scan fluorescence intensity profiles (green/red channels) across the nuclear region of the cell shown in (**A**). **(C)** Co-condensation of EMB1579 with CDKC;2 *in vitro*. Oregon Green 488-labeled CDKC;2 was incubated with rhodamine-labeled EMB1579 under crowding conditions. Scale bars, 5 μm. **(D)** Line-scan fluorescence intensity profiles (green/red channels) across the condensate shown in (**C**).

To characterize the interaction, we sought to identify the specific region(s) of CDKC;2 responsible for binding. IUPred2A predicted that the extended C-terminus (residues 368-513) of CDKC;2 constitutes an intrinsically disordered region (IDR), designated CDKC;2-IDR. CDKC;2 also contains a core kinase domain comprising CDKC;2-N (N-lobe, residues 1-124) and CDKC;2-C (C-lobe, residues 125-367), encompassing the ATP-binding site, cyclin interface, and substrate-binding sites (Bosken et al. 2014) (Figure S5A).

We first assessed these regions for interaction with EMB1579 using a yeast two-hybrid (Y2H) system. Surprisingly, full-length CDKC;2 did not show detectable interaction; however, both CDKC;2-N and CDKC;2-IDR exhibited strong interactions (Figure S5B). This suggests that the full-length interaction is weak or transient and may be masked by intramolecular contacts in the Y2H context. To further delineate the interface, we tested two truncation mutants: CDKC;2ΔN (lacking the N-lobe) and CDKC;2ΔIDR (lacking the IDR). CDKC;2ΔN retained interaction with EMB1579, whereas CDKC;2ΔIDR showed no detectable interaction (Figure S5B). These results indicate that the IDR is necessary and presumably sufficient for the EMB1579-CDKC;2 interaction, whereas the N-lobe contributes only marginally.

Given the weak, likely electrostatic nature of this interaction, we hypothesized that EMB1579 and CDKC;2 may undergo co-condensation. *In vitro* phase separation assays supported this: EMB1579 and CDKC;2 colocalized within the same condensates (Figure 3C, D), and CDKC;2 exhibited a heterogeneous distribution within EMB1579 condensates, aggregating into small foci (Figure 3C). Interestingly, both CDKC;1 and CDKC;2 could partition non-selectively into the same EMB1579 condensates (Figure S6), providing an explanation for their redundant roles in Pol II CTD phosphorylation.

### EMB1579 Concentrates CDKC;2 in a Concentration-Dependent Manner

Given that EMB1579 regulates CDKC;2 function, the two proteins co-condense *in vitro* and colocalize in the nucleus, and EMB1579 phase separation is required for its physiological function (Zhang et al. 2020b), we therefore asked whether EMB1579 LLPS is specifically required for its role in regulating CDKC;2. Co-sedimentation assays confirmed that CDKC;2 partitioning into condensed phases increased with higher EMB1579 concentrations under LLPS conditions (Figure S7). This concentration-dependent effect was also confirmed by fluorescence microscopy: compared with CDKC;2 alone, CDKC;2 condensates were dramatically enhanced by increasing EMB1579 concentrations (Figure 4A). Pearson’s correlation coefficient analysis indicated that increasing EMB1579 concentrations elevated the proportion of CDKC;2 within EMB1579 condensates (Figure 4B).

**Figure 4.**
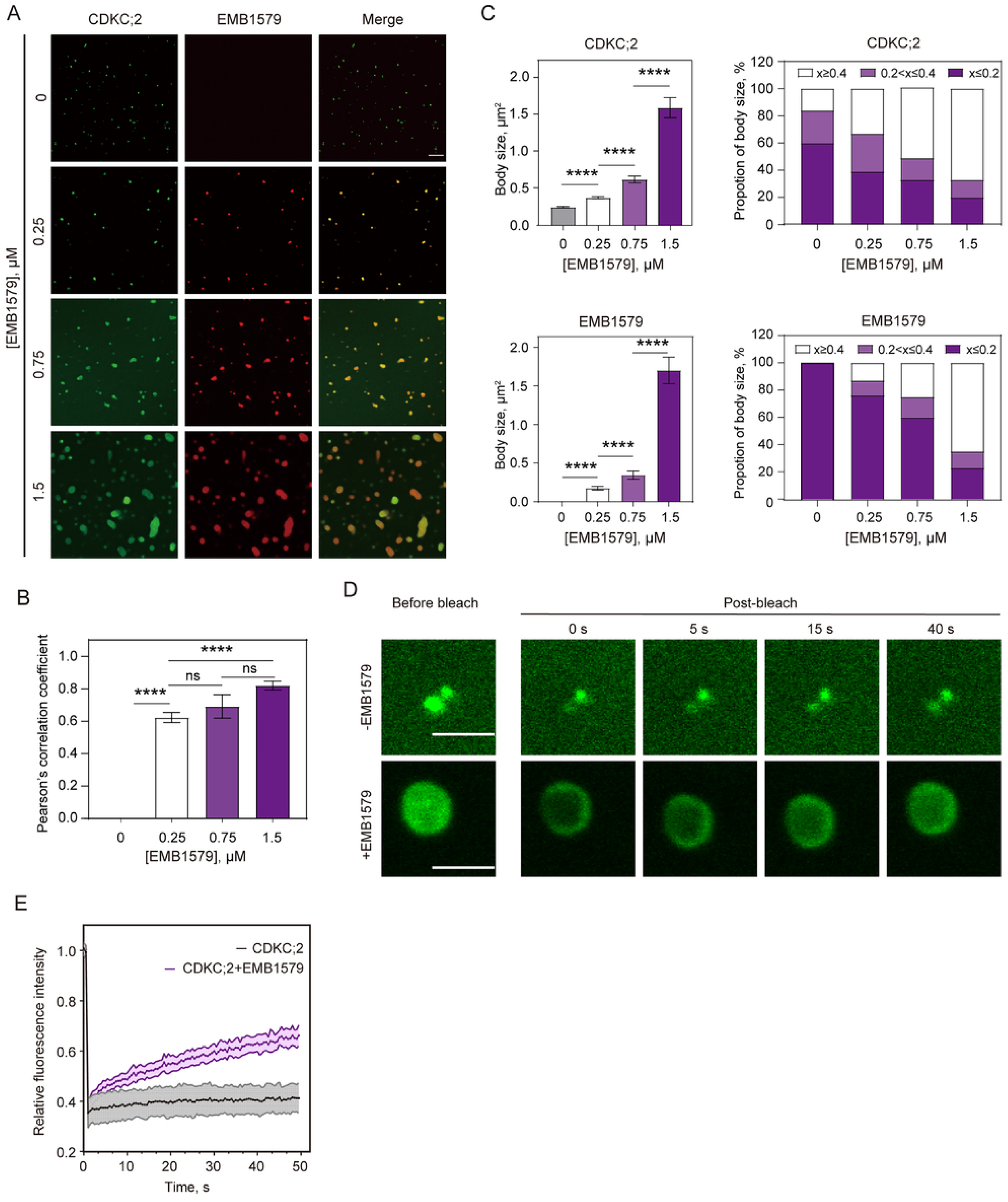
EMB1579 Recruits and Concentrates CDKC;2 to Form High-order Condensates *in Vitro*. **(A)** Representative images of Oregon Green 488-CDKC;2 mixed with 0 μM, 0.25 μM, 0.75 μM, or 1.5 μM rhodamine-EMB1579 in phase separation reaction buffer. Scale bar, 5 μm. **(B)** Pearson’s correlation coefficients for the co-localization analysis of CDKC;2 and EMB1579 in (**A**). The data are means ± SEM; n=5. Asterisks indicate significant differences between the samples with different concentrations of EMB1579 (two-tailed paired Student’s *t* test, \*\*\*\**P* ≤ 0.0001, ns, not significant). **(C)** Quantitative analyses of body size of CDKC;2 in (**A**). The condensates size of CDKC;2 are divided into three clusters of different body size (μm^2^). The proportion of body size refers to the ratio of the number of droplets in three clusters to the total droplets number. The data are means ± SEM. Asterisks indicate significant differences between the samples with different concentrations of EMB1579 (two-tailed paired Student’s *t* test, \*\*\*\**P* ≤ 0.0001). (**D** and **E**) FRAP analysis of Oregon Green 488-CDKC;2 proteins recovery rate. In panel D, 0 s represents the starting time point of recovery after photobleaching, and the rates of fluorescence recovery was plotted every 0.5 s. The rates of fluorescence recovery before photobleaching are set to 1.0. The data are means ± SD; n = 10.

Quantitative analysis revealed that the mean condensate size of CDKC;2 increased significantly with rising EMB1579 concentrations. At a constant CDKC;2 concentration, the mean area of CDKC;2 in the presence of 2 μM EMB1579 (1.6 μm²) was approximately 6.4-fold larger than in the absence of EMB1579 (0.25 μm²). The proportion of condensates exceeding 0.4 μm² increased accordingly, suggesting that EMB1579 promotes the fusion of CDKC;2 into higher-order assemblies (Figure 4C). FRAP imaging showed that CDKC;2 fluorescence recovered from the condensate surface toward the center, resulting in a well-distributed pattern, indicating that CDKC;2 aggregation occurs after recruitment by EMB1579 (Figure 4D).

Quantification of fluorescence recovery further revealed rapid recovery of CDKC;2 within EMB1579 condensates, in contrast to minimal recovery in CDKC;2-only droplets (Figure 4E), indicating that the fluidity of CDKC;2 increases upon recruitment into EMB1579 condensates. In summary, EMB1579 recruits CDKC;2 and concentrates it into restricted spatial regions within EMB1579 condensates in a concentration-dependent manner.

### LLPS of EMB1579 Is Required for Enhancing CDKC;2 Kinase Activity

To test whether EMB1579 phase separation is required for this functional interaction, we analyzed CDKC;2 distribution in protoplasts expressing EMB1579^ΔRED^, a mutant lacking the RED domain (residues 351-428) that disrupts phase separation (Zhang et al. 2020b). Unlike wild-type EMB1579, EMB1579^ΔRED^ induced a more diffuse CDKC;2 distribution (Figure 5A, B), quantified by a 40% reduction in the coefficient of variation (CV) of CDKC;2 fluorescence (Figure 5C). This indicates that EMB1579 phase separation promotes CDKC;2 condensation *in vivo*. Despite maintaining protein interactions with CDKC;2 domains (e.g., N-terminus and IDR; Figure S8), EMB1579^ΔRED^ failed to restore Pol II CTD phosphorylation in *emb1579* mutants (Figure 5D), demonstrating that phase separation, rather than protein interaction alone, is required for EMB1579 to regulate CDKC;2-mediated CTD phosphorylation. Consistent with its failure to fully rescue *FLC* transcription defects (Zhang et al. 2020b), proper EMB1579 phase separation is critical for both Pol II CTD phosphorylation and *FLC* transcriptional regulation.

**Figure 5.**
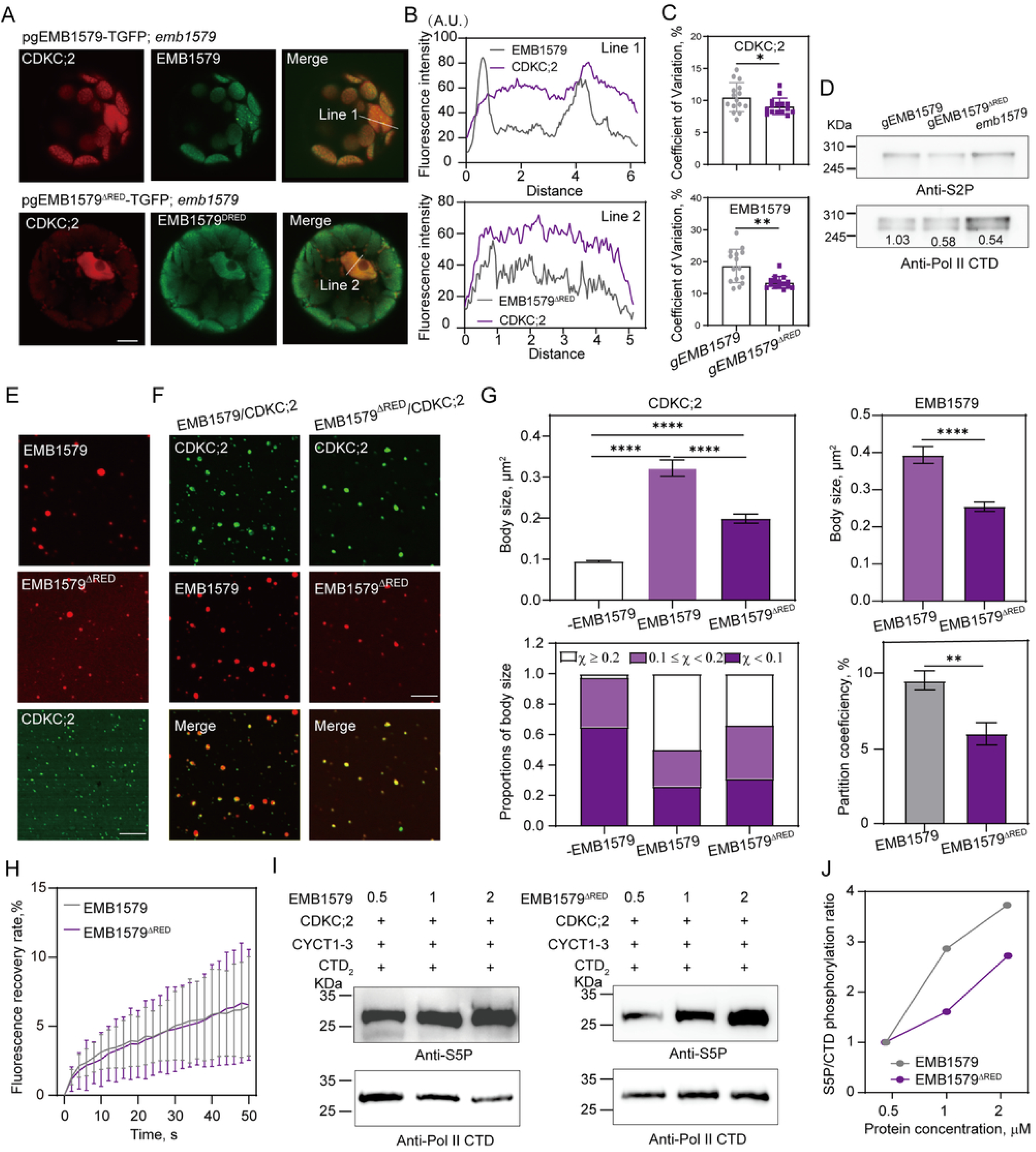
Phase Separation Is Essential for EMB1579 to Recruit CDKC;2 and Enhance Pol II CTD Phosphorylation. (**A**, **B**) Subcellular localization of RFP-CDKC;2 in *Arabidopsis* mesophyll protoplasts of gEMB1579 or gEMB1579^DRED^ (**A**). Line-intensity profiles of the green and red channels across two randomly selected lines through the nuclei are illustrated on the fluorescent images (**B**). Scale bar: 5 μm. **(C)** The coefficient of variation of fluorescence of RFP (CDKC;2) or GFP (EMB1579 or EMB1579^DRED^) in gEMB1579 or gEMB1579^DRED^. The data are means ± SEM; n = 15. Asterisks indicate significant differences (two-tailed paired Student’s *t* test, \**P* ≤ 0.05; \*\**P* ≤ 0.01). **(D)** RNA pol II Ser2 phosphorylation ratio in different genetic backgrounds. Western blots of protein extracted from 7-day-old seedlings. Membranes were probed with antibodies against CTD Ser2 P Pol II (3E10) or total Pol II (8WG16). Ser2 phosphorylation level normalized to total CTD are shown relative to that in gEMB1579 set at 1.0. **(E)** Representative images of 1 μM Oregon Green 488-CDKC;2 or 2 μM rhodamine-EMB1579, EMB1579^DRED^ in phase separation reaction buffer alone. Scale bar, 5 μm. **(F)** Representative images of Oregon Green 488-CDKC;2 incubated with rhodamine-EMB1579 or rhodamine-EMB1579^DRED^. Scale bar, 5 μm. **(G)** Top panel: quantitative analyses of body size of CDKC;2 or EMB1579 in panel B. Bottom panel: the proportion of CDKC;2 condensates of different areas and partition coefficient. **(H)** FRAP analysis of Oregon Green 488-CDKC;2 proteins recovery rate. 0 s represents the starting time point of recovery after photobleaching, and the rates of fluorescence recovery was plotted every 0.5 s. (**I**, **J**) Phosphorylation of RNA pol Ⅱ with different concentration of EMB1579 or EMB1579^DRED^. In vitro, purified EGFP-CDKC;2, CYCT1-3-YFP and GST-CTD_2_ were incubated with different concentrations of EMB1579 or EMB1579^DRED^ in phosphorylation reaction buffer. Membranes were probed with antibodies against CTD Ser5 P Pol II (D9N5I) or total Pol II (8WG16). Phosphorylation of RNA pol Ⅱ CTD Ser5 normalized to total CTD with 0.5 µM EMB1579 or EMB1579^DRED^ set at 1.0.

To dissect how EMB1579 phase separation regulates CDKC;2 kinase function, we compared the condensate properties of wild-type (WT) EMB1579 and EMB1579^ΔRED^, as well as their effects on CDKC;2 kinase activity. Under molecular crowding conditions (100 mg/mL PEG3350), EMB1579^ΔRED^ formed smaller, more numerous condensates compared to WT EMB1579 (Figure 5E), indicating impaired phase separation. CDKC;2 condensates co-assembled with WT EMB1579 exhibited a 1.5-fold larger mean area and a 2.3-fold higher proportion of droplets >0.2 μm² than those with EMB1579^ΔRED^ (Figure 5F, G). Both variants increased condensed-phase partitioning of CDKC;2 relative to CDKC;2 alone, but WT EMB1579 showed superior recruitment efficiency (Figure 5F, G). FRAP revealed comparable dynamics of CDKC;2 within WT and mutant EMB1579 condensates (Figure 5H), indicating that the RED domain modulates phase separation scale without altering CDKC;2-EMB1579 interaction strength. Functionally, both variants enhanced CDKC;2-mediated phosphorylation of Pol II CTD at Ser5, but WT EMB1579 was significantly more effective (Figure 5I, J). The reduced activity of EMB1579^ΔRED^ correlates with its diminished condensate size and recruitment capacity, establishing that proper EMB1579 phase separation is critical for optimal CDKC;2 kinase function.

### EMB1579 Condensates Recruit the P-TEFb Complex and Promote Pol II CTD Phosphorylation

In plants, CDKC;2, the ortholog of human CDK9, associates with Cyclin T partners (e.g., CYCT1-3, CYCT1-4, CYCT1-5), and CYCT1-3 specifically activates CDKC;2 to phosphorylate Pol II *in vitro* (Barroco et al. 2003; Li et al. 2014). To investigate whether EMB1579 spatially interacts with the P-TEFb complex, we performed co-visualization of Pol II, CYCT1-3, and EMB1579 in cells. Confocal microscopy revealed that while most Pol II distributed diffusely in the nucleus, a subset formed discrete foci that partially overlapped with EMB1579 condensates (Figure 6A). Furthermore, when CYCT1-3-RFP was co-expressed with EMB1579-TGFP in protoplasts, CYCT1-3 foci colocalized with EMB1579 condensates (Figure 6B, C). These observations suggest that EMB1579 condensates spatially overlap with CDKC;2-containing P-TEFb complexes *in vivo*.

**Figure 6.**
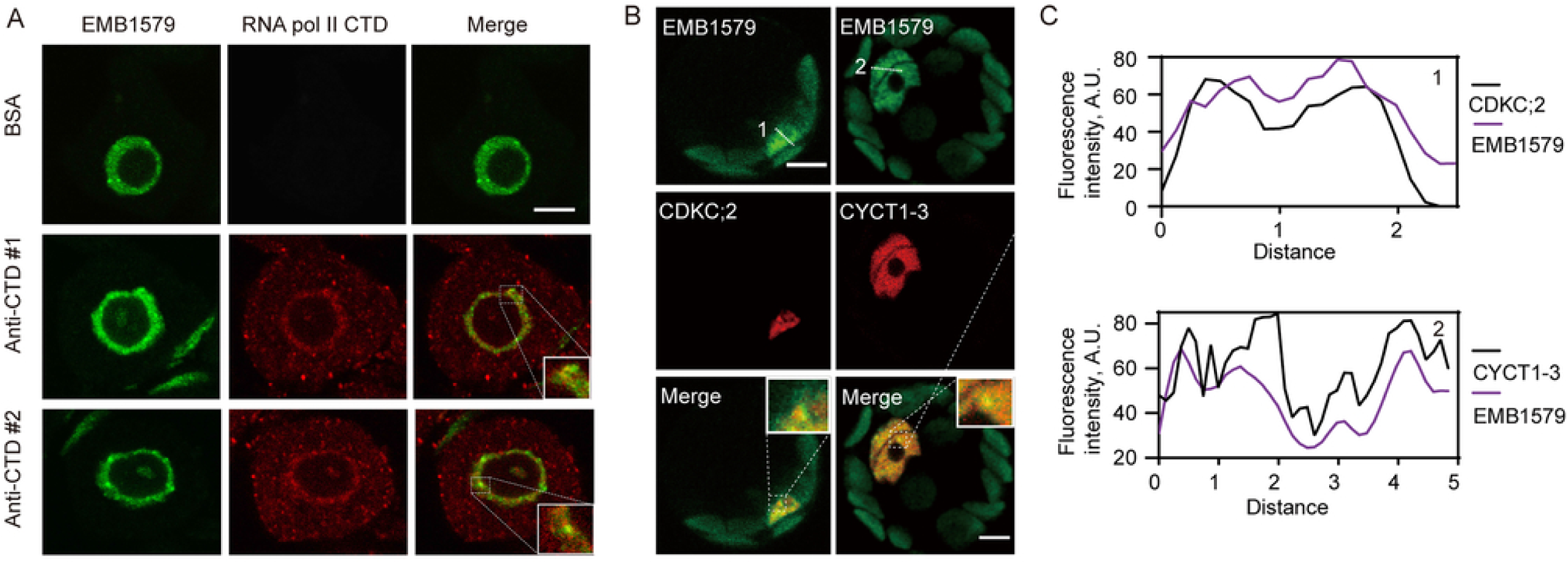
Partial Co-localization of P-TEFb Complex and RNA Pol II within EMB1579 Condensates. **(A)** Immunofluorescence analysis of RNA polymerase II (Pol II) localization in 4-day-old *gEMB1579* seedling roots. Membranes were probed with anti-Pol II antibody (8WG16), with BSA serving as a negative control to exclude non-specific binding. Scale bar, 5 μm. **(B)** Subcellular localization of RFP-tagged P-TEFb components (CDKC;2 and CYCT1-3) in *Arabidopsis* mesophyll protoplasts expressing *gEMB1579*. Line-scan fluorescence intensity profiles (green/red channels) across nuclear regions (illustrated on fluorescent images) demonstrate co-localization patterns. Scale bar, 5 μm. **(C)** Line-scan fluorescence intensity analysis. A line segment was manually drawn across the target subcellular region, and the fluorescence intensity along this line was measured. The plotted profile shows fluorescence intensity per individual pixel (y-axis) as a function of distance along the line (x-axis). Line 1 and line 2 indicate the colocalization of CDKC;2 and EMB1579, and colocalization of CYCT1-3 and EMB1579, respectively.

To determine whether EMB1579 directly recruits P-TEFb and Pol II *in vitro*, we first demonstrated that EMB1579, CDKC;2, CYCT1-3, and the CTD_2_ peptide co-assemble within the same biomolecular condensate (Figure 7A, B). We then examined whether EMB1579 influences P-TEFb complex assembly. Under crowding conditions (100 mg/mL PEG3350), CDKC;2, CYCT1-3, and CTD_2_ formed irregular, aggregated condensates that failed to fuse into larger structures (Figure 7C), indicating that additional regulators are required for proper P-TEFb architecture. Strikingly, the addition of EMB1579 promoted the formation of well-defined, spherical P-TEFb condensates, with a significant increase in both size and fluorescence intensity (Figure 7D). These data demonstrate that EMB1579 actively recruits and concentrates P-TEFb components into functional condensates, thereby enhancing their capacity to phosphorylate Pol II CTD at Ser2 (Figure 7E).

**Figure 7.**
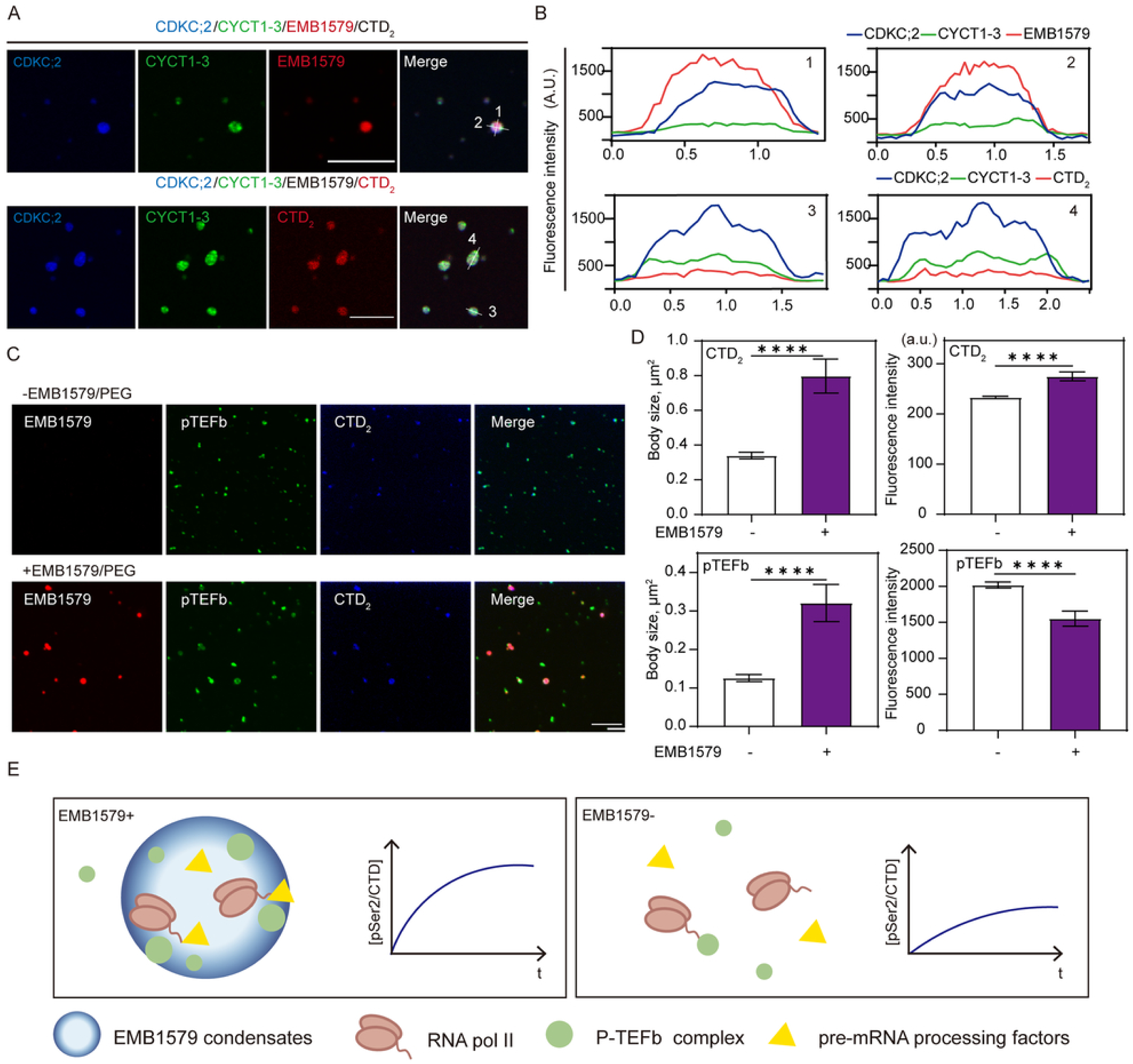
P-TEFb Complex and RNA Pol II Incorporate into EMB1579 Condensates. **(A)** Confocal images of phase-separated condensates formed by rhodamine-EMB1579 (top) or rhodamine-GST-CTD2 (bottom) with iFluor 405-CDKC;2, CYCT1-3-YFP, and GST-CTD2 in buffer containing 100 mg/mL PEG3350. Scale bars, 5 μm. **(B)** Line-scan fluorescence intensity profiles across selected nuclear regions (as indicated in panel A) showing co-localization of EMB1579, CDKC;2, CYCT1-3, and CTD2. **(C)** Confocal images of EGFP-CDKC;2, CYCT1-3, and iFluor 405-GST-CTD2 with/without rhodamine-EMB1579 in 100 mg/mL PEG3350. Scale bars, 5 μm. **(D)** Quantitative analysis of P-TEFb (CYCT1-3/*CDKC;2*) and CTD2 condensate size distributions and fluorescence intensity from panel C. Data are mean ± SEM. Asterisks denote significant differences (two-tailed paired Student’s t-test: ****P ≤ 0.0001). **(E)** Schematic model describing the role of EMB1579 in forming condensates in promoting the function of pTEFb complex in promoting transcription via phosphorylating pol II CTD.

## Discussion

Pol II CTD phosphorylation orchestrates transcription and pre-mRNA processing (Bird, Zorio, and Bentley 2004; Corden 2013; Coudreuse et al. 2010; Jeronimo, Bataille, and Robert 2013), processes fundamental to plant growth and development. CDKC;2 is the primary Ser2 kinase in Arabidopsis, and its depletion causes pleiotropic defects including circadian rhythm disruption, delayed flowering, and immunodeficiency (Cui et al. 2007; Li et al. 2014; Wang et al. 2014; Zhao et al. 2017a; Uehara et al. 2022). This study identifies EMB1579 as a novel regulator of CDKC;2 activity through LLPS, establishing a mechanism in which EMB1579 condensates act as reaction hubs that recruit and concentrate CDKC;2, CYCT1-3, and CTD peptides, thereby enhancing CDKC;2 kinase activity toward CTD substrates (Figure 7E). This phase-separated microenvironment facilitates efficient Pol II CTD phosphorylation, providing critical insights into transcription regulation in plants.

A confluence of multiple convergent lines of evidence identifies EMB1579 as an upstream regulator of CDKC;2 in the pathway controlling Pol II CTD phosphorylation and global transcriptional regulation. Our core results show that EMB1579 depletion markedly lowers *in vivo* Ser2 phosphorylation (Figure 2A). Recombinant EMB1579 enhances CDKC;2 kinase activity toward Pol II CTD substrates *in vitro* (Figure 2B, C), an effect fully dependent on EMB1579’s phase separation activity both *in vitro* (Figure 5I, J) and *in vivo* (Figure 5D). Genetically, combined loss of EMB1579 and CDKC;2 synergistically upregulates *FLC* transcript levels to cause delayed flowering (Figure 2F-H), while *emb1579cdkc;2* double mutants exhibit more severe developmental defects and greater reductions in Pol II CTD phosphorylation than either single mutant (Figure 2F-J), confirming the hierarchical epistatic relationship between the two factors.

The more severe phosphorylation defect in *emb1579cdkc;2* double mutants compared to *cdkc;2* single mutants (Figure 2I, J) is likely attributed to EMB1579’s regulatory role over CDKC;1, a functionally redundant paralog of CDKC;2. Consistent with this hypothesis, both CDKC;1 and CDKC;2 co-localize with EMB1579 within biomolecular condensates (Figure S6). These findings delineate a hierarchical regulatory network in which EMB1579 serves as a master regulator, coordinating the recruitment and activation of CDKC kinases, including both CDKC;1 and CDKC;2, to precisely control Pol II CTD phosphorylation dynamics.

Disruption of EMB1579’s LLPS significantly reduces Pol II CTD phosphorylation (Figures 5D), elevates *FLC* transcription, and delays flowering (Zhang et al. 2020b), demonstrating its critical regulatory function. EMB1579 interacts with CDKC;2 at least part through its IDR, and its LLPS dynamics are modulated by local concentration thresholds and phase transition boundaries (Zhang et al. 2020b). This regulatory mechanism mirrors mammalian systems, where P-TEFb is recruited into transcriptional elongation condensates to hyperphosphorylate the Pol II CTD and enhance elongation rates (Lu et al. 2018), underscoring the evolutionary conservation of phase separation in transcriptional control. Notably, CDKC;2 can form condensates independently of EMB1579 (Figure 5E), suggesting context-dependent regulatory mechanisms. Future studies should dissect the spatiotemporal conditions under which EMB1579 modulates CDKC;2 activity at specific genomic loci.

CTD phosphorylation regulates Pol II interactions during transcription cycles, governing promoter-proximal pausing and co-transcriptional processes such as splicing, polyadenylation, and termination. Aberrant phosphorylation alters transcription speed, generating alternative transcripts and impacting transcriptome composition, a mechanism conserved across eukaryotes but with plant-specific adaptations. For instance, Pol II with hyperphosphorylated CTD interacts with RNA splicing factors during elongation, forming splicing condensates that facilitate pre-mRNA processing (Peng, Li, and Xu 2020; Guo et al. 2019). EMB1579 may bridge transcription and splicing by orchestrating the transition from elongation to splicing condensates, consistent with its proposed roles in splicing regulation (Zhang et al. 2020b).

Unexpectedly, *FLC* transcription is more profoundly disrupted in *emb1579* mutants than in *cdkc;2* mutants (Figure 1E), indicating that EMB1579 regulates *FLC* expression through additional targets beyond CDKC;2. This aligns with prior studies showing that EMB1579 employs multifaceted mechanisms to control *FLC* transcription. For instance, EMB1579 has been implicated in PRC2-mediated H3K27me3 deposition (Zhang et al. 2020b), and a physical interaction between EMB1579 and FCA has been identified (Fang et al. 2020), suggesting potential modulation of FCA-dependent pathways. However, direct evidence confirming EMB1579’s functional influence on FCA remains to be established. These findings underscore the necessity of comprehensive interactome analyses to elucidate the full scope of EMB1579’s role in transcriptional regulation.

In summary, this study advances understanding of phase separation in plant transcription regulation and positions EMB1579 as a key coordinator of Pol II-centered processes. By elucidating the interplay between LLPS and Pol II CTD phosphorylation, it provides a framework for investigating plant-specific adaptations in transcriptional control and highlights the need for comprehensive interactome analyses to uncover additional regulatory partners.

## Material and Method

### Plant Materials and Growth Conditions

Arabidopsis thaliana ecotype Columbia-0 (Col-0, wild type) and two previously characterized mutant alleles, *cdkc;2-2* (SALK_029546C) and *emb1579-3* (SALK_007142), were employed in this study, as described in prior work (Zhao et al. 2017b; Zhang et al. 2020a). For simplicity, the *cdkc;2-2* and *emb1579-3* lines are hereafter referred to as *cdkc;2* and *emb1579*, respectively. The double mutant *emb1579 cdkc;2* was generated via genetic crossing between homozygous single-mutant lines, and successful homozygosity was verified by PCR using primer pairs flanking the T-DNA insertion sites (detailed in Table S1). Two previously generated transgenic lines, gEMB1579 and gEMB1579^DRED^ (Zhang et al. 2020b), were also included for subsequent functional analyses.

Seeds were surface-sterilized, stratified at 4°C for 3-5 days, and plated on ½-strength Murashige and Skoog (1/2 MS) medium supplemented with 1% sucrose, 1% agar, and species-specific antibiotics for selection. Plates were incubated at 22°C under a 16-h-light/8-h-dark cycle for 7 days in a tissue culture chamber. Seedlings were then transplanted to soil and grown in a greenhouse under identical photoperiod conditions.

### Plasmid Construction

To assess the interaction between EMB1579 and CDKC;2, the *CDKC;2* coding sequence was PCR-amplified with primers introducing EcoRI/SacI sites and cloned into pGADT7 (bait vector) via restriction digestion, generating pGADT7-CDKC;2. Truncated *CDKC;2* variants were constructed using identical cloning strategies. The pGBKT7-EMB1579 and pGBKT7-EMB1579^DRED^ prey vectors were prepared as previously described (Zhang et al. 2020b).

For *in vivo* visualization of CDKC;2 and CYCT1-3 subcellular localization, mRFP-tagged fusion constructs were generated. The *CDKC;2* and *CYCT1-3* coding sequences were amplified via PCR using gene-specific primers (PEZS-CDKC;2-F/R, PEZS-CYCT1-3-F/R) (Table S1) and inserted into linearized PEZS-mRFP-NL (via homologous recombination) to yield PEZS-mRFP-CDKC;2-NL and PEZS-mRFP-CYCT1-3-NL. Linearization of PEZS-mRFP-NL was achieved by PCR amplification with PEZS-F/R primers (Table S1).

Full-length *CDKC;2* was amplified from *Arabidopsis* seedling cDNA using pET28a-CDKC;2-F/R primers and cloned into EcoRI-digested pET28a via homologous recombination, generating pET28a-CDKC;2. The pColdI-EGFP-CDKC;2 construct was created by inserting *CDKC;2* (amplified with pColdI-CDKC;2-F/R) into EcoRI/SalI-digested pColdI-GFP (Zhang et al. 2020b). For CYCT1-3-YFP production, *YFP* was first cloned into EcoRI/SalI-digested pET28a to generate pET28a-YFP. Subsequently, *CYCT1-3* (amplified with pET28a-CYCT1-3-F/R (Table S1)) was inserted into BamHI/EcoRI-digested pET28a-YFP via homologous recombination, yielding pET28a-CYCT1-3-YFP. The pGEX-CTD2 construct was prepared as previously reported (Cui et al. 2007).

### Generation of Transgenic Plants

To silence *CDKC;1* expression in wild-type (Col-0) or *emb1579cdkc;2* double mutants, a 300-bp fragment from the 3’ untranslated region (3’ UTR) of *CDKC;1* was PCR-amplified using primers *CDKC;1* RNAi-F (XbaI/AscI sites) and *CDKC;1* RNAi-R (BamHI/SwaI sites; Table S1). Target selection followed established criteria (Cui et al. 2007). The amplified fragment was cloned into the pFGC5941 vector via restriction digestion. Resultant constructs were transformed into *Arabidopsis* using *Agrobacterium*-mediated floral dip (Clough and Bent 1998). Basta-resistant T1 transgenic plants were selected and confirmed by PCR.

To generate *pCDKC;2:tdTomato-CDKC;2* and *pEMB1579:gEMB1579-TGFP* complemented lines, homozygous *pEMB1579:gEMB1579-TGFP; emb1579cdkc;2* plants were first produced by crossing *gEMB1579* (Zhang et al. 2020b) with *cdkc;2* mutants. The *proCDKC;2* promoter was amplified from genomic DNA using Pro-CDKC;2-F/R primers, while *gCDKC;2* was amplified with gCDKC;2-F/R primers (Table S1). The pFGC5941-tdTomato vector and *proCDKC;2* fragment were digested with StuI/SwaI to create an intermediate construct (pFGC5941-*proCDKC;2*-tdTomato). *gCDKC;2* was then inserted into this intermediate via AscI/BamHI digestion, yielding pFGC5941-*proCDKC;2*-tdTomato-*gCDKC;2*. This construct was transformed into *pEMB1579:gEMB1579-TGFP; emb1579cdkc;2* plants via floral dip. Basta-resistant T1 plants were selected and validated by fluorescence microscopy.

### Recombinant Protein Expression and Purification

For EGFP-CDKC;2, CYCT1-3-YFP, and CDKC;2 production, plasmids pColdI-EGFP-CDKC;2, pET28a-CYCT1-3-YFP, or pET28a-CDKC;2 were transformed into *Escherichia coli* Rosetta (DE3) cells. Cultures (2 L LB medium) were grown at 37°C to OD₆₀₀ ≈ 0.6. Protein expression was induced with 0.2 mM IPTG, followed by overnight incubation at 15°C. Cells were lysed in lysis buffer (25 mM Tris-HCl pH 8.0, 250 mM KCl, 5 mM imidazole, 1 mM PMSF, lysozyme) via sonication. Lysates were centrifuged (12,000 × *g*, 4°C, 1 h), and supernatants were incubated with Ni-NTA resin. Resin was washed with buffer containing 30 mM imidazole, and proteins were eluted in 600 mM imidazole buffer. Purified proteins were dialyzed against storage buffer (25 mM Tris-HCl pH 8.0, 500 mM KCl) prior to freezing.

GST-CTD2 was purified using glutathione agarose beads. The pGEX-CTD2 construct (validated by Sanger sequencing) was transformed into *E. coli* BL21 (DE3). Cells were lysed in PBS (140 mM NaCl, 2.7 mM KCl, 10 mM Na₂HPO₄, 1.8 mM KH₂PO₄), centrifuged (12,000 × *g*, 4°C, 1 h), and supernatants were incubated with glutathione agarose. The column was washed with PBS until no protein was detected in flow-through. GST-CTD2 was eluted in PBS containing 50 mM glutathione, dialyzed against preservation buffer (25 mM Tris-HCl pH 8.0, 250 mM KCl), and stored at −80°C. EMB1579 and EMB1579ᴰᴿᴱᴰ proteins were purified as previously described (Zhang et al. 2020b).

### Transient Expression in Arabidopsis Mesophyll Protoplasts

Mesophyll protoplasts were isolated from *gEMB1579* or *gEMB1579^DRED^* transgenic plants (Zhang et al. 2020b) using the PEG-calcium-mediated transformation protocol (Yoo, Cho, and Sheen 2007). Plasmids pEZS-NL-RFP-CDKC;2 or RFP-CYCT1-3 (10-20 μg) were transfected into protoplasts (2 × 10⁴ cells/mL) and incubated for 8**-**12 hours at 22°C in the dark. Subcellular localization was analyzed using an Olympus FV1200MPE laser scanning confocal microscope. mRFP fluorescence was detected with a 561 nm laser line (emission 570-620 nm), while TGFP fluorescence was visualized with a 488 nm laser (emission 500-550 nm). Images were processed using Fiji/ImageJ for contrast adjustment and co-localization analysis.

### In Vitro Kinase Assays

Kinase assays were performed with modifications to the protocol described by Uehara *et al*. (2022). EGFP-CDKC;2 and EMB1579/EMB1579ᴰᴿᴱᴰ proteins were dialyzed into kinase reaction buffer (20 mM HEPES-KOH pH 8.0, 8 mM MgCl₂, 0.5% glycerol, 1 mM DTT) prior to use. Reaction mixtures contained 2.5 μM GST-CTD_2_, 0.25 μM CYCT1-3-YFP, 50 μM MG132, protease inhibitor cocktail (1×), 5 mM ATP, and varying concentrations of CDKC;2 and EMB1579 proteins in a final volume of 30 μL. Reactions were incubated at room temperature (22-25°C) for 90 min with gentle mixing every 10 min to ensure homogeneity. The reaction was terminated by adding 5 μL of 5× SDS-PAGE loading buffer and heating at 95°C for 10 min. Samples were resolved on 12.5% SDS-PAGE gels and visualized by Coomassie staining or immunoblotting as required.

### Pol II Phosphorylation In Vivo Analysis

Wild-type *Arabidopsis* seedlings (7-day-old, n=20 per replicate) grown on ½ MS plates were harvested and flash-frozen in liquid nitrogen. For inhibitor treatments, seedlings (n=15 per condition) were soaked in BML-259 (0.1-10 μM) for 30 min at 22°C in a tissue culture chamber prior to liquid nitrogen freezing. Total protein was extracted using 1.5× Laemmli buffer (supplemented with 8 M urea, 1× protease inhibitor cocktail, and 1× phosphatase inhibitors) at 10 μL per mg fresh weight. Extracts were centrifuged (20,000 × *g*, 45 min, 4°C) to remove debris, and supernatants were heated at 95°C for 10 min.

Protein samples (equivalent to 10 mg fresh weight) were resolved on 6% SDS-PAGE gels (Morimoto et al. 2013) and transferred to PVDF membranes. Pol II was detected using anti-RNA polymerase II monoclonal antibody (8WG16, 1:2000; ab817, Abcam). Ser2-phosphorylated CTD was visualized with Ser2P CTD antibody (3E10, 1:1000; Active Motif). Immunoblots were developed using ECL Prime reagent and imaged with a ChemiDoc MP system. Band intensities were quantified using Fiji/ImageJ, with normalization to total Pol II signal.

### Real-Time RT-PCR

Seven-day-old *Arabidopsis* seedlings grown under a 16-h-light/8-h-dark photoperiod were harvested and flash-frozen in liquid nitrogen. Total RNA was extracted using the Eastep® Super Total RNA Extraction Kit (LS1040, Promega) following the manufacturer’s protocol. RNA (5 μg) was reverse-transcribed into cDNA using the SuperScript™ IV First-Strand Synthesis System (M1705, Invitrogen). cDNA was diluted (1:10) and used as template for quantitative PCR performed on an ABI 7500Fast Real-Time PCR System with RealStar SYBR Mixture (A301, Genstar). Gene-specific primers (Table S1) were designed for target genes. Relative expression was calculated using the ΔΔCt method, with normalization to reference genes. Data are presented as means ± SEM from three independent biological replicates.

### Yeast Two-Hybrid (Y2H) Assay

To assess protein interactions, pGADT7 (bait) and pGBKT7 (prey) plasmids were co-transformed into *Saccharomyces cerevisiae* Gold strain competent cells using the LiAc/SS-DNA/PEG method (Clontech). Transformed yeast were plated on synthetic dropout (SD) medium lacking leucine and tryptophan (SD/-Leu/-Trp) and incubated at 30°C for 2–3 days to select for plasmid co-expression. Resultant colonies were then streaked onto SD medium lacking histidine, leucine, and tryptophan (SD/-His/-Leu/-Trp) supplemented with 3-amino-1, 2, 4-triazole (3-AT; 1-5 mM) to suppress background His3 expression. Plates were incubated at 30°C for 3 days, and growth was scored as a positive interaction.

### In Vitro Phase Separation Assay

CDKC;2 and EMB1579/EMB1579ᴰᴿᴱᴰ proteins were pre-clarified by centrifugation (12,000 × *g*, 4°C, 5 min). Proteins were labeled with Oregon Green 488 (CDKC;2) or rhodamine (EMB1579 variants) in dialysis buffer (25 mM Tris-HCl pH 8.0, 100 mM KCl, 2 mM DTT) at 4°C in the dark for 2 h. Labeled proteins were dialyzed into LLPS buffer (25 mM Tris-HCl pH 8.0, 100 mM KCl, 2 mM DTT) overnight at 4°C. Reaction mixtures (10 μL final volume) contained 3 μM Oregon Green 488-CDKC;2 and variable concentrations of rhodamine-EMB1579 (0.5-3 μM). Mixtures were injected into flow chambers with Poly-L-lysine-coated coverslips. Condensate formation was observed at 22°C using an Olympus FV1200MPE confocal microscope with 488 nm (Oregon Green) and 561 nm (rhodamine) excitation. Partition coefficients (ratio of protein in condensates vs. dilute phase) were calculated as described (King and Petry 2020). Dynamic exchange within CDKC;2-EMB1579 condensates was analyzed by FRAP. Bleaching was performed using a 488 nm laser (100% power, 2 s pulse). Fluorescence recovery was monitored at 5-s intervals for 2 min and quantified using Fiji/ImageJ. Recovery rates were calculated as previously described (Zhang et al. 2020b).

### Low-Speed Co-Sedimentation Assay

To assess protein-protein interactions via phase separation, CDKC;2 and EMB1579/EMB1579ᴰᴿᴱᴰ were pre-clarified by centrifugation (12,000 × *g*, 4°C, 5 min) and dialyzed overnight into LLPS buffer (25 mM Tris-HCl pH 8.0, 100 mM KCl, 2 mM DTT) at 4°C. Reaction mixtures (40 μL final volume) contained 2.5 μM CDKC;2, variable concentrations of EMB1579 variants (0.5-3 μM), and 100 mg/mL PEG3350 to induce phase separation. Mixtures were centrifuged at 13,600 × *g* for 30 min at 4°C to separate soluble (supernatant) and condensed (pellet) fractions. The pellet was gently resuspended in an equal volume of LLPS buffer to match the supernatant volume. Both fractions were analyzed by 10% SDS-PAGE with Coomassie blue staining. Band intensities were quantified using Fiji/ImageJ. The partition coefficient (pellet/supernatant ratio) was calculated to assess interaction strength.

### Immunofluorescence Assay

Root tips (0.5-1 cm) from 4-day-old *Arabidopsis* seedlings were excised and fixed in 4% (w/v) paraformaldehyde in PME buffer (50 mM PIPES, 5 mM MgCl₂, 5 mM EGTA, pH 6.9) supplemented with 0.05% (v/v) Triton X-100 for 1-1.5 h at 22°C. Fixed tissues were washed twice with PME buffer (5 min each) and once with PBS (pH 7.0) to remove residual fixative. Root tips were digested with 1% (w/v) cellulase R-10 and 0.02% (w/v) pectolase in PBS for 10 min at 22°C to soften cell walls. Tissues were then washed three times with PBS (5 min each) to remove enzymatic residues. Poly-L-lysine-coated coverslips were prepared by incubating in 0.1 mg/mL poly-L-lysine solution for 10 min, followed by air-drying. Digested root tips were gently transferred onto coverslips, and two coverslips were pressed together to disperse cells. Samples were flash-frozen in liquid nitrogen and carefully separated to generate intact cell monolayers. Pol II was detected using anti-RNA polymerase II monoclonal antibody (8WG16, 1:500; ab817, Abcam) diluted in PBS containing 1% BSA and 0.1% Triton X-100. Primary antibodies were incubated overnight at 4°C. Secondary detection employed Alexa Fluor 568-conjugated anti-mouse IgG (1:1000; A11004, Invitrogen) for 2 h at 22°C. Coverslips were mounted in ProLong Gold antifade reagent (P36930, Invitrogen) for imaging. Samples were visualized using an Olympus FV1200MPE confocal microscope with 561 nm laser excitation (emission 570-620 nm). Negative controls (no primary antibody or non-specific IgG) were included to validate staining specificity. Image analysis was performed using Fiji/ImageJ, with fluorescence intensity quantified in defined regions of interest.

### BML-259 Treatments

*Arabidopsis* seedlings (7-day-old) grown on ½ MS medium under a 16-h-light/8-h-dark photoperiod were treated with BML-259 (5 or 20 μM) or an equivalent volume of DMSO (mock control) in liquid ½ MS medium. Treatments were performed at 22°C for 30 min under constant light to maintain photoperiod consistency. Following treatment, seedlings (n=3 biological replicates per condition) were flash-frozen in liquid nitrogen for subsequent RNA or protein extraction. DMSO-treated seedlings served as negative controls to account for solvent effects. All treatments were performed in triplicate to ensure statistical robustness.

## Acknowledgements

We thank the members of the Huang Lab for their insightful discussions. We are also thankful to the Arabidopsis Biological Resource Center and the Nottingham Arabidopsis Stock Centre for generously providing T-DNA insertion lines. This work was supported by grants from the National Key R&D Program of China (2022YFA1303400) and the National Natural Science Foundation of China (32570809). Declaration of generative AI and AI-assisted technologies in the writing process: The authors acknowledge the use of DeepSeek for assistance with grammatical corrections and improved phrasing.

## Author contributions

XZ developed the experimental methods; XZ and YD performed the majority of the experiments, with conceptual input from SH. XZ, YD, YZ and PH analyzed the data with conceptual input from SH. SH conceived the study, and supervised the experimental work. ZX and SH wrote the manuscript. All authors have carefully read and approved the manuscript.

## Data availability

The data that supports the findings of this study are available within the article and its Supporting Information (Figs S1–S8).

---

**Figure S1. CDKC;1 and CDKC;2 are enriched in EMB1579 pull-down fraction**

**(A)** Functional enrichment analysis of EMB1579-associated biological processes. The top 10 enriched terms, derived from Metascape analysis of previous study data (Zhang et al. 2020b), are shown, with “regulation of transcription by RNA polymerase II” highlighted in red. Data were obtained from Metascape (https://metascape.org/).

**(B)** Scores of CDKC family proteins CDKC;1 and CDKC;2 from the EMB1579 pull-down coupled mass spectrometry analysis, and their corresponding rankings in the “RNA polymerase II-mediated transcriptional regulation” category.

**Figure S2. Expression patterns of *CDKC;1* (AT5G10270) and *CDKC;2* (AT5G64960).**

**(A)** Overall expression patterns of *CDKC;1* (AT5G10270) and *CDKC;2* (AT5G64960) across developmental stages, from seed and seedling to the mature plant.

(**B**–**F**) Detailed comparison of the two homologous genes in specific organs, including the embryo (B), shoot apex (C), leaf (D), silique (E), and the entire rosette after the transition to flowering (F). Colors indicate expression levels in RPKM (reads per kilobase per million mapped reads) based on Araport 11 RNA-seq data. Images were generated with the AtGenExpress eFP at bar.utoronto.ca/eplant (Waese et al. 2017) and adapted for this figure. RSE, root mean square error.

**Figure S3. Loss of CDKC;2 delays flowering**

**(A)** Gene structure of *CDKC;2* (AT5G64960). The red arrow indicates the T-DNA insertion site in the eighth exon of the *cdkc;2* mutant (SALK_029546). Blue arrows denote 5’ and 3’ UTR, respectively. Black squares represent exons.

**(B)** Genotyping of *cdkc;2* and genomically complemented plants by a three-primer PCR assay. gCDKC;2 designates the complementation line in which the *cdkc;2* mutant was transformed with the genomic sequence of *CDKC;2*.

**(C)** Relative expression of *CDKC;2* in *cdkc;2* and the complemented seedlings.

**(D)** Late flowering phenotypes of *cdkc;2* plants compared with wild type and gCDKC;2. Scale bar, 2 cm.

**Figure S4. BML259 treatment reduces Pol II phosphorylation and elevates *FLC* transcription in both WT and *emb1579*.**

**(A)** *In vivo* Pol II phosphorylation in 7-day-old wild-type and *emb1579* seedlings treated with BML259 (0, 5, or 20 μM) or equivalent volume of DMSO in liquid medium for 30 min at room temperature. Total protein was extracted and analyzed by immunoblotting, with total Pol II CTD as loading control.

**(B)** Phosphorylation levels were normalized to DMSO-treated wild-type or *emb1579*, set to 1. Left: representative immunoblots; right: quantification (mean ± SD, n = 3).

**(C)** *FLC* transcript levels measured by RT-qPCR from the same samples, normalized to *TUB8*. DMSO-treated wild-type was set to 1. Data shown as mean ± SD (n = 3).

**Figure S5. EMB1579 interacts with IDR of CDKC;2**

EMB1579 interacts with IDR of CDKC;2 as determined by yeast two-hybrid assays. The top panel shows a schematic representation of CDKC;2 domain structure. IDR, intrinsic disordered region. -WH, SD-Leu-Trp medium. -WLH, SD-His-Leu-Trp medium.

**Figure S6. Both CDKC;1 and CDKC;2 can cocondense with EMB1579**

CDKC;1, CDKC;2, and EMB1579 were labeled with Oregon Green, rhodamine, and iFluor 405, respectively, mixed in vitro, and immediately imaged. Horizontal panels show condensation of each protein across different conditions; vertical panels represent different protein combinations. Rightmost column: 5× magnified images of condensates formed by CDKC;1, CDKC;2, and EMB1579. Scale bars, 5 μm.

**Figure S7. CDKC;2 cosediments with EMB1579 condensates in Vitro**

Low-speed co-sedimentation analysis of CDKC;2 and EMB1579 partitioning between dilute (supernatant, S) and condensed (pellet, P) phases. Asterisks indicate EMB1579 (140-180 kDa) and CDKC;2 (45-60 kDa) bands. Numbers below CDKC;2 bands denote the ratio of supernatant/pellet proteins to total input.

**Figure S8. Loss of RED Motif Does Not Disrupt EMB1579-CDKC;2 Interaction**

**(A)** Schematic representation of the RED motif localization within the EMB1579 protein domain architecture. The RED motif (red box) is positioned in the intrinsically disordered region (IDR).

**(B)** Yeast two-hybrid assay confirming interaction between EMB1579^ΔRED^ (RED motif deletion mutant) and the IDR of CDKC;2. Top panel: Growth assays on selective media (-WH: SD-Leu-Trp medium for plasmid selection; -WLH: SD-His-Leu-Trp medium for interaction detection) demonstrate specific interaction. IDR, intrinsic disordered region.

